# Beige fat appearance during lipoatrophy reveals trade-offs between metabolic homeostasis and female fertility

**DOI:** 10.1101/2025.04.21.649835

**Authors:** Elizabeth S. Anaya, William Dion, Pradip K. Saha, Aaron R. Cox, Evelyn de Groot, Avery Ahmed, Jessica B. Felix, Bokai Zhu, Stephanie A. Pangas, Sean M. Hartig

## Abstract

White adipose tissue (WAT) meets the energetic demands required for estrous cycle regularity, ovulation, and reproductive success in women. Here, we report WAT loss and subsequent female-specific compensatory beige fat recruitment impact metabolic and reproductive outcomes in mice. Female mice that undergo progressive lipoatrophy (‘fat-less’) displayed disrupted estrous cycles, reduced ovarian reserve, and subfertility. These effects were attributable to an accumulation of thermogenic beige fat cells in residual subcutaneous WAT depots and greater energy expenditure. Conversely, high-fat diet lowered energy expenditure and rescued estrous cycle regularity among ‘fat-less’ mice compared to littermate controls, despite profound insulin resistance and metabolic dysfunction. Together, these findings provide evidence of trade-offs between the maintenance of metabolic homeostasis and reproduction, where increased energy expenditure and beige fat cell emergence compensate for fat loss but occur at the expense of fecundity in female mice.

## Introduction

White adipose tissue (WAT) responds to the body’s metabolic demands, storing excess energy as triglycerides during nutrient overload and releasing free fatty acids in times of nutrient deficit (Ghaben and Scherer 2019). WAT has been recognized for its importance in reproduction beyond traditional roles as an energy reservoir. In females, body composition is a significant risk factor for infertility, with excessive adipose tissue seen in overweight or obese individuals increasing risks of ovulatory disorders and decreasing success in fertility treatments (Zhu et al. 2022). Additionally, reduced fertility is also found in individuals with diminished adipose tissue, such as athletes, women with lipodystrophies, or eating disorders, including anorexia and bulimia nervosa (Zhu et al. 2022; Boutari et al. 2020). These observations further demonstrate that maintaining quality adipose tissue is required to sustain metabolic homeostasis and reproductive success.

Considerable evidence indicates the amount of thermogenic, beige adipocytes within subcutaneous WAT (sWAT) depots corresponds with insulin sensitivity and protection against the co-morbidities of obesity and aging (Chouchani and Kajimura 2019). On the contrary, some heterogeneous disorders characterized by fat atrophy couple thermogenesis in WAT with systemic lipid and carbohydrate metabolism defects (Agarwal et al. 2021; Srinivasa et al. 2019; Gallego-Escuredo et al. 2013; Pellegrini et al. 2019; Béréziat et al. 2011; Rodríguez de la Concepción et al. 2004). It is still unclear whether beige fat recruitment can be translated into effective treatments for obesity. However, the implications of beige fat activity on female fertility remain largely unknown. It is known that females accumulate adipose tissue differently than males (Mauvais-Jarvis 2024) and exhibit a greater capacity for thermogenic activity within brown and white fat depots, often attributed to higher estrogen levels (Santos et al. 2018; Cypess et al. 2009; Kim et al. 2016). Yet, the consequences of beige fat cell recruitment on female reproduction, including cycle regularity, have not been explored.

Here, we explore sex-specific metabolic adaptations to fat loss and female reproductive outcomes using a lipodystrophic mouse model (*Ubc9^flox/flox^*; *Adipoq*-Cre)(Cox et al. 2021). Previously, we showed that the small ubiquitin-like modifier (SUMO) E2-conjugating enzyme *Ubc9* is important for WAT expansion (Cox et al. 2021). In this study, we initially focused on the long-term metabolic outcomes resulting from a female-specific increase in sWAT beige fat resulting from WAT loss. However, we surprisingly observed that *Ubc9^flox/flox^*; *Adipoq*-Cre mice (*Ubc9^fKO^*, ‘fat-less’) female mice also have reduced fertility. To these points, our studies reveal significant consequences of sex-specific beige fat recruitment, whereby compensatory beige fat activity diverts energy to thermogenesis instead of being available for reproductive function.

## RESULTS

### ‘Fat-less’ mice lose WAT in a sex-specific and age-dependent manner

During our previous investigations of ‘fat-less’ *Ubc9^fKO^*mice, we noticed sexually dimorphic metabolic phenotypes, including less subcutaneous WAT (sWAT) inflammation and less lipid accumulation in the liver of the females compared to the males (Cox et al. 2021). To investigate these observations further, we measured subcutaneous (sWAT) and visceral (vWAT) tissue weights across five time points in male and female ‘fat-less’ mice. Noticeable fat loss in *Ubc9^fKO^*mice occurred post-puberty in both sexes but varied between depots and across time points. During the progressive fat loss, female mice maintained more vWAT (Figures 1A and 1B) but lost sWAT earlier than their male counterparts (Figures 1C and 1D). At the 6-month-old timepoint, when male and female *Ubc9^fKO^* mice have significantly less adipose tissue in both depots compared to controls, they also display differences in circulating serum hormone levels. Serum insulin is drastically increased in *Ubc9^fKO^* males, but this increase is not observed in the females, as their insulin levels are comparable to littermate controls (Figure 1E). Additionally, WAT loss in both sexes reduced serum adiponectin and leptin levels, yet to a lesser degree in females (Figure 1E). These findings built upon previous studies (Cox et al. 2021) and demonstrated sex-specific differences in response to the metabolic stress of progressive fat loss.

**Figure 1.**
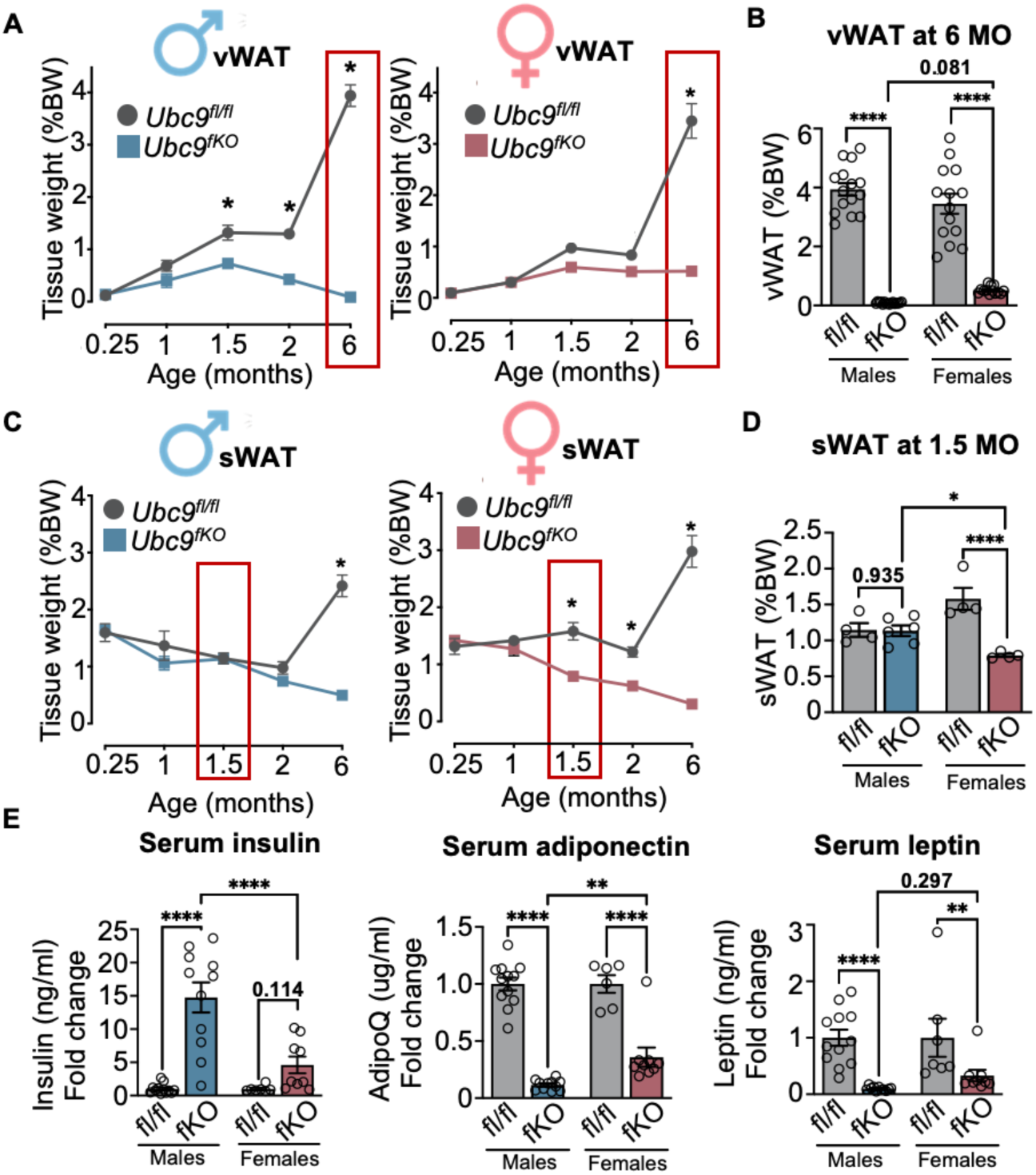
‘Fat-less’ mice lose WAT in a sex-specific and age-dependent manner. (A) Visceral white adipose tissue (vWAT) weights from necropsy over time from male (blue) and female (pink) *Ubc9^fKO^* mice and *Ubc9^fl/fl^* littermate controls (n=3-16/group). Red boxes represent sex differences in vWAT loss. Data are mean ± SEM. *p<0.05, by two-way ANOVA followed by Fisher’s LSD Test. (B) Sex comparison of vWAT weights at 6 months of age (n=7-16/group). Tissue weights are shown as percent body weight (BW). (C) Subcutaneous white adipose tissue (sWAT) weights from necropsy over time (n=3-10). Red boxes represent sex differences in sWAT loss. Data are mean ± SEM. *p<0.05 by two-way ANOVA followed by Fisher’s LSD Test. (D) Sex comparison of sWAT weights at 1.5 months of age (n=4,5/group). Tissue weights are shown as percent body weight (BW). (E) Male and female comparison of serum insulin, adiponectin (adipoQ), and leptin levels measured ad libitum at 6 months of age (n=7-12/group). Data are mean ± SEM. *p<0.05, **p<0.01, ***p<0.001, ****p<0.0001 by two-way ANOVA followed by Fisher’s LSD post-hoc test.

### Female *Ubc9^fKO^* remain insulin sensitive despite loss in adipose tissue mass and quality

We next characterized adipose tissue morphology changes from two months of age, when fat loss begins in female ‘fat-less’ mice, as well as in aged 11-month-old mice. At two months of age, *Ubc9^fKO^*females display smaller and more multilocular-appearing adipocytes in WAT depots compared to littermate controls, which were most apparent in the sWAT depot (Figure 2A). Interestingly, two-month-old *Ubc9^fKO^* female mice weighed more than their control counterparts, likely due to increased lean mass (Figure 2B). Age also strongly affected adipose tissue morphology among genotypes. Consistent with previous observations in six-month-old *Ubc9^fKO^* mice (Cox et al. 2021), hypertrophic and inflamed adipocytes were observed at 11 months of age, with only a few small remnant adipocytes. As a consequence, we found depleted levels of the adipokines leptin and adiponectin synthesized from the residual adipocytes in aged 11-month-old *Ubc9^fKO^* mice but not at two months (Figure 2C). Smaller adipocytes in younger mice contributed to higher blood serum free fatty acids, while lipoatrophy later in life demonstrably reversed effect sizes observed in two-month-old female *Ubc9^fKO^* mice, with no difference in triglycerides at either age (Figure 2D). Serum cholesterol levels increased by almost 50% in older mice, which coincided with drastically heightened insulin levels, which were unchanged in two-month-old mice (Figure 2E).

**Figure 2.**
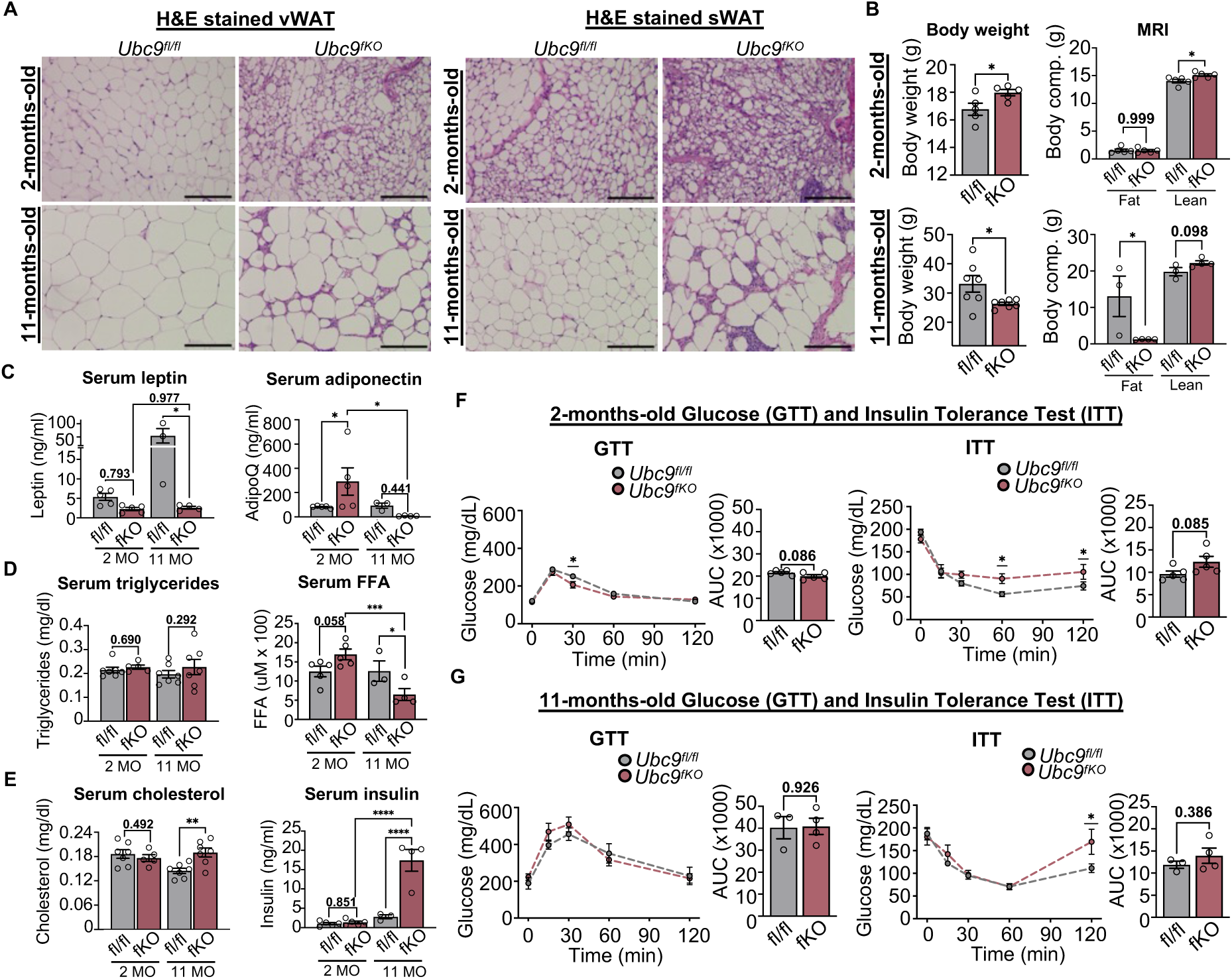
Female *Ubc9^fKO^* remain insulin sensitive despite loss in adipose tissue mass and quality. (A) H/E-staining of visceral (vWAT) and subcutaneous (sWAT) white adipose tissue from 2-month-old and 11-month-old female *Ubc9^fKO^* mice and *Ubc9^fl/fl^* littermate controls (n=3,4/group). All images were taken at 20x. Scale bar set at 100µm. (B) 2-month-old (n=5/group) and 11-month-old (n=3,4/group) body weight and body composition measured by Echo MRI, shown as percentage of body weight. (C) Serum adipokines leptin and adiponectin (AdipoQ) from ad libitum fed mice (n=3-7/group). (D) Serum triglycerides and free fatty acids (FFA) levels measured ad libitum (n=3-7/group). (E) Serum cholesterol and insulin (n=3-7/group). Data are mean ± SEM. *p<0.05, **p<0.01 by two-way ANOVA followed by Fisher’s LSD post-hoc test for serum measurements and body composition and unpaired two-tailed student’s *t*-test for body weight. (F) Glucose tolerance test (GTT) and insulin tolerance test (ITT) with corresponding area-under-curve (AUC) measurements at 2-months (n=5). (G) GTT, ITT, and corresponding area-under-curve (AUC) measurements at 11 months. Data are mean ± SEM. *p<0.05 by repeated measures ANOVA for GTT and ITT and unpaired two-tailed student’s *t*-test for AUC.

*Ubc9^fKO^* mice and littermate controls were then subjected to glucose (GTT) and insulin tolerance tests (ITT) to determine how high insulin levels impacted insulin and glucose tolerance. Surprisingly, both age groups remained similarly glucose tolerant and mostly insulin sensitive (Figures 2F and 2G), which did not demonstrably change with age. These data identified progressive dyslipidemia and altered adipokine secretion in female *Ubc9^fKO^* mice, which largely occur because of WAT atrophy. Yet, young and old female *Ubc9^fKO^* mice responded to glucose and insulin similarly to their littermates and, thus, for the most part, are metabolically healthy despite the metabolic stress associated with fat loss.

### Young *Ubc9*^fKO^ female mice recruit beige fat to compensate for WAT atrophy

To characterize whole-body energy balance and substrate utilization near the onset of fat loss, we placed two-month-old female *Ubc9*^fKO^ mice in CLAMS cages and explored phenotypes after correcting for lean body mass using CalR (Mina et al. 2018). Indirect calorimetry revealed an unexpected increase in energy expenditure (Figure 3A) and a shift in substrate utilization in female *Ubc9^fKO^* mice compared to *Ubc9^fl/fl^* littermate controls, as indicated by a lower respiratory exchange ratio (RER) in the active (dark) phase (Figure 3B). No differences in food intake were observed (Figure 3C).

**Figure 3.**
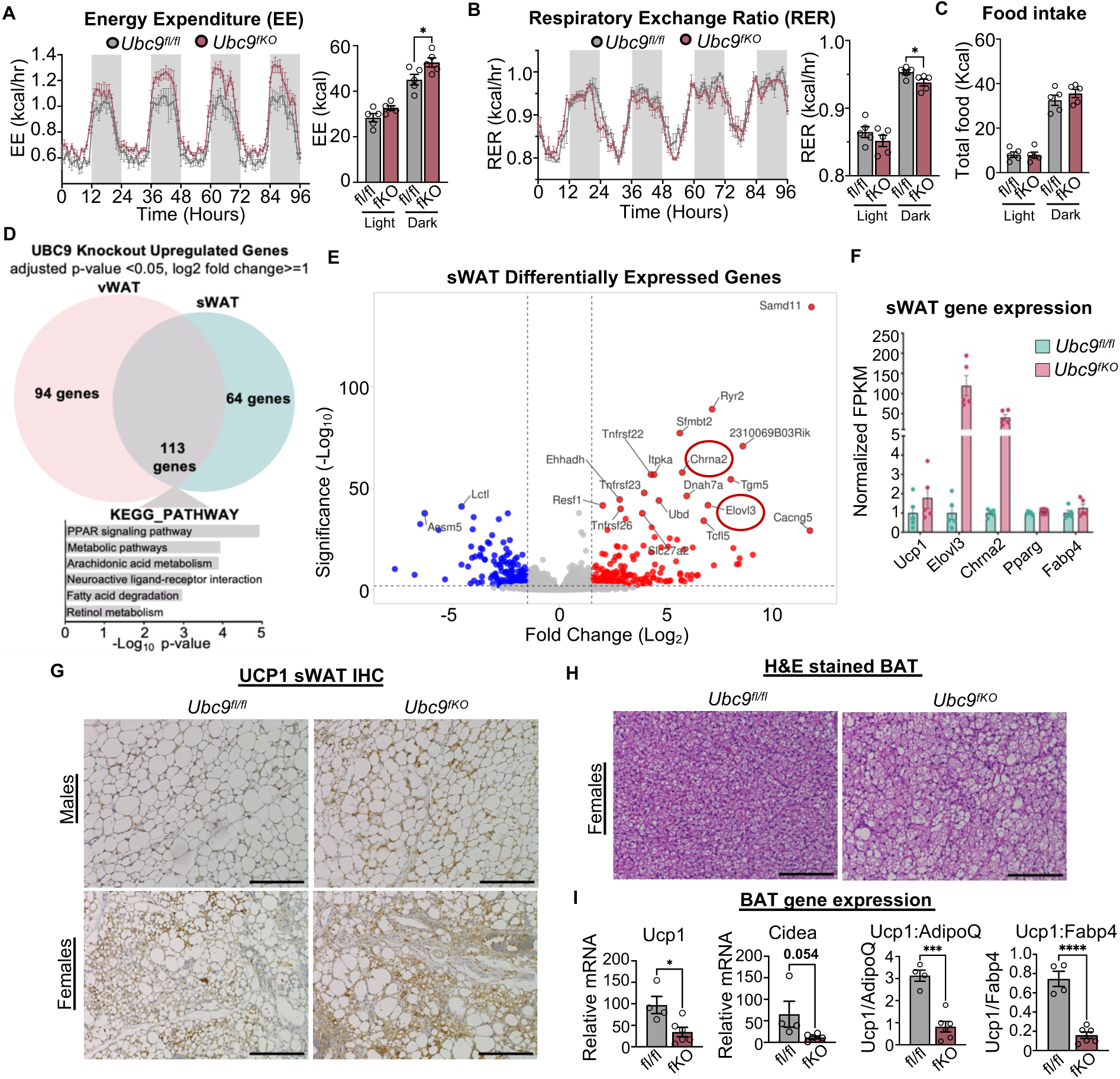
Young *Ubc9^fKO^* female mice recruit beige fat to compensate for WAT atrophy. 2-month-old *Ubc9^fKO^* and *Ubc9^fl/fl^* littermate controls were individually housed and monitored in CLAMS cages during a 96 h period. Statistical analysis performed by ANCOVA with lean body mass as a covariate for body mass-dependent variables (EE, and food intake) and ANOVA for mass-independent variables (RER) using CalR web-based tool (n=5/group). Data are mean ± SEM. *p<0.05, **p<0.01, ***p<0.001. (A) Energy expenditure (EE). (B) Respiratory exchange ratio (RER). (C) Average total food intake in light and dark phase. (D) RNAseq analysis of WAT from 2-month-old mice with adjusted p-value <0.05, log2 fold change>=1 and KEGG-Pathway analysis (n=5/group). (E) sWAT differentially expressed genes highlighting highly expressed genes (red circles). (F) RNA expression shown as normalized FPKM in sWAT. (G) Immunohistochemistry staining of UCP1 in 2-month-old male and female *Ubc9^fKO^* mice and *Ubc9^fl/fl^* littermate controls mice and corresponding measured expression shown as percent area of stain. All images were taken at 20x. Scale bar set at 100µm (n=3/group). (H) H&E stained brown adipose tissue (BAT) from 2-month-old *Ubc9^fKO^* and *Ubc9^fl/fl^* littermate controls. Images taken at 20x. Scale bar set at 100µm (n=3/group). (I) Relative mRNA of thermogenic genes from BAT (n=3,4/group). Data are mean ± SEM. *p<0.05, ***p<0.001 by unpaired two-tailed student’s *t*-test.

Next, we performed bulk RNA-Seq of WAT depots to determine broad gene expression changes occurring with whole-body metabolic changes among female *Ubc9^fKO^* mice compared to *Ubc9^fl/fl^* littermate controls. These efforts uncovered clear signatures that explain the impacts of *Ubc9* knockout in the vWAT and sWAT (Figure 3D). Kyoto Encyclopedia of Genes and Genomes (KEGG) analysis identified that *Ubc9^fKO^* vWAT and sWAT shared higher expression of genes enriched in pathways known to favor adipocyte functions, including PPAR signaling and lipid metabolism. Lowered RER is associated with enhanced thermogenic activity and accumulation of beige fat (Petruzzelli et al. 2014; Guo et al. 2022). As such, we were not surprised to find thermogenic genes *Elovl3* and *Chrna2* to be two of the most upregulated genes in the sWAT (Figure 3E). *Elovl3* strongly responds to the recruitment of brown adipose tissue in response to cold (Westerberg et al. 2006) and serves as a marker of beige adipocytes (Jakobsson et al. 2005; Chen et al. 2019) along with *Chrna2,* which can act in both adrenergic-dependent and independent pathways (Jun et al. 2020). White fat genes *Pparg* and *Fabp4* remain unchanged, while *Ucp1,* an important marker of thermogenesis and WAT browning, was marginally increased in sWAT from *Ubc9^fKO^* mice (Figure 3F).

To build confidence that *Ubc9^fKO^* female mice accumulated more beige fat, sWAT from two-month-old female mice was analyzed by immunohistochemistry for UCP1 expression and compared to littermate controls and males. We observed morphological and UCP1 expression changes in sWAT across sexes and genotypes. sWAT excised from female *Ubc9^fKO^* mice contained a greater abundance of smaller, multilocular adipocytes compared to males and littermate controls, indicative of more beige fat cells. Immunohistochemistry further confirmed these cells were UCP1+ adipocytes in *Ubc9^fKO^*female mice (Figure 3G). *Ubc9^fKO^* males appeared to have slightly higher UCP1 expression than control males. However, a substantial increase in UCP1 levels was observable in female knockouts compared to all other groups. Beige fat appearance is sexually dimorphic (Kim et al. 2016) and can occur differently in males and females in response to dietary and environmental stressors. As such, sWAT beige adipocyte emergence in female *Ubc9^fKO^* may confer adaptive responses to compensate for the loss of *Adipoq*+ fat reserves.

Some studies argue that the overall contribution of beige fat to metabolic activity is considerably lower than that of brown adipose tissue (BAT) (Labbé et al. 2016). As such, we also analyzed the brown adipose tissue (BAT) from these mice to determine whether the enhanced energy expenditure could be explained by increased BAT thermogenesis. Instead, BAT excised from two-month-old *Ubc9^fKO^* females contained more unilocular adipocytes, apparent in H&E-stained histological sections (Figure 3H). Additionally, reduced expression of BAT genes (*Ucp1*, *Cidea*) further confirmed more whitening in *Ubc9^fKO^* females (Figure 3I) and fails to explain the higher energy expenditure seen at this age. Our findings indicate that beige fat recruitment during WAT loss contributed to systemic energy expenditure and increased lipid oxidation.

### Estrous cycle irregularity and subfertility precede lipoatrophy in female mice

The impacts of beige fat on female reproduction remain uncharacterized. Given the increased energy expenditure associated with beige adipocytes within intact sWAT observed in female *Ubc9^fKO^* mice, we next investigated whether estrous cycle regularity and fertility were affected. 30-day estrous cycle monitoring in two-month-old female *Ubc9^fKO^*mice revealed irregular cycles (Figure 4A), characterized by a reduction in metestrus days, an increase in days spent in estrus, and an overall greater proportion of estrus days per month (Figure 4B). Because *Ubc9^fKO^*displayed abnormal estrous cycles, we conducted a six-month-long fertility assessment in which sexually mature eight-week-old *Ubc9^fKO^* and *Ubc9^fl/fl^* controls were continuously mated with age-matched sexually mature wild-type (WT) males. In doing so, we saw a significant reduction in total pups born over time from *Ubc9^fKO^* dams (Figure 4C). The overall litter size of *Ubc9^fKO^* dams decreased by two pups on average (Figure 4D). Additionally, in the last two months of the fertility study, the time between litters significantly increased in *Ubc9^fKO^*mice, averaging over 40 days (Figure 4E). Interestingly, after birth, a significant number of pups did not survive past weaning age (Figure 4F). Of note, premature death in which pups do not survive to weaning is seen in other lipodystrophic models (Wang et al. 2013) and is often attributed to pup fat loss. However, *Ubc9^fKO^* does not generate a germline mutation, and *Ubc9^fKO^* females were bred to reproductively young WT males; thus, offspring are heterozygous for the *Ubc9* floxed allele and not knockouts. Pup loss was likely a result of early onset WAT dysfunction and inadequate nutritional supply from fat knockout dams.

**Figure 4.**
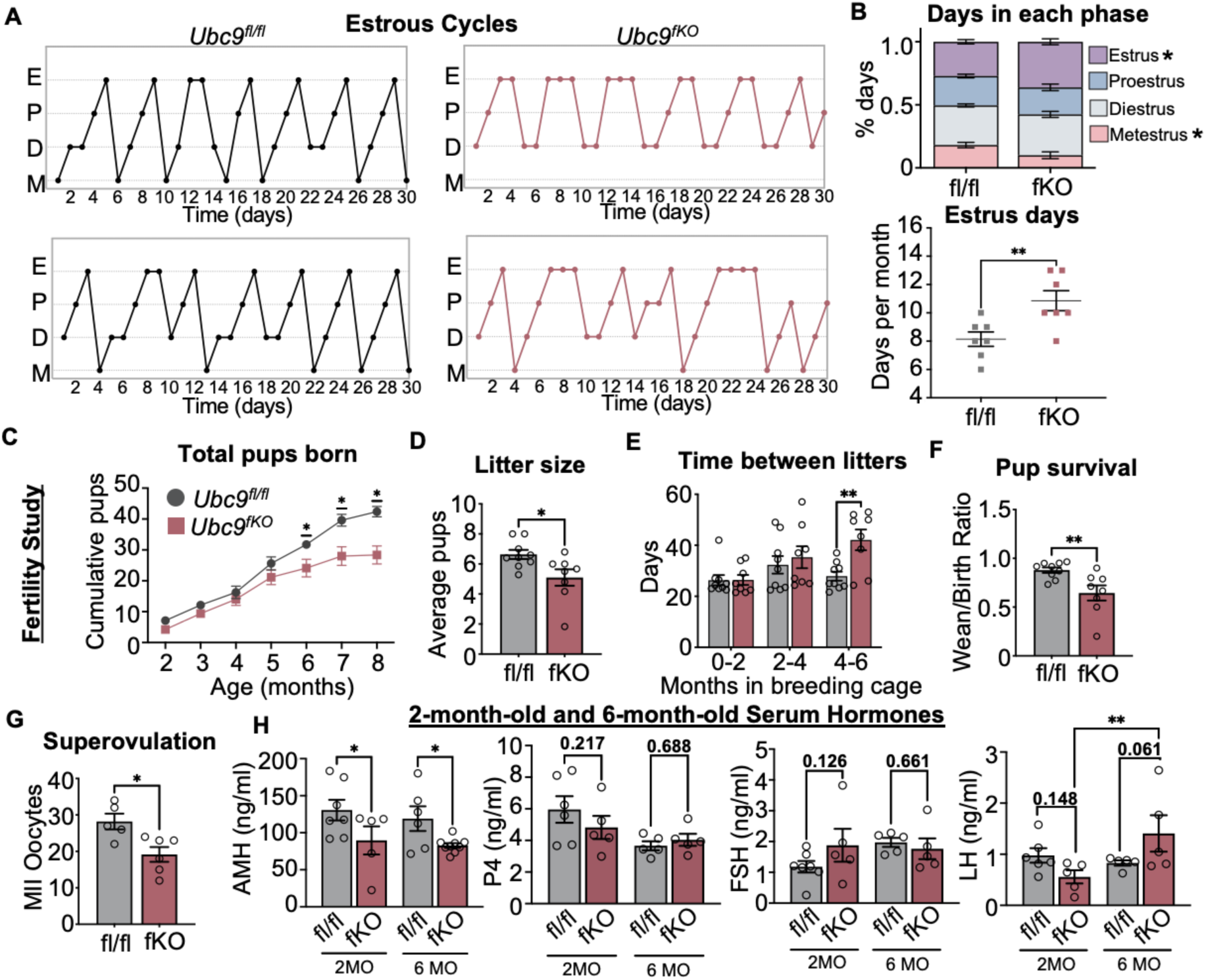
Estrous cycle irregularity and subfertility precede lipoatrophy in female mice. (A) Representative 30-day estrous cycle monitoring in 2-month-old *Ubc9^fKO^* and littermate controls (n=7/group). (B) Percent days in each phase and number of days spent in estrus over a 30-day period (n=7/group). Data are mean ± SEM. *p<0.05, by two-way ANOVA followed by Fisher’s LSD post-hoc test. (C) Total number of pups born per female over a 6-month-long fertility study mated to WT mice starting at 2 months of age (n=9,8/group). Data are mean ± SEM. *p<0.05, by repeated measures ANOVA. (D) Overall average litter size per female over 6-month period. (E) Time in days between litters per female separated into 2-month-long periods. Data are mean ± SEM. **p<0.01, by repeated measures ANOVA. (F) Pup survival measured by as a ratio between number of pups weaned and born per female. (G) Number of MII oocytes ovulated from superovulation experiment using 5-week-old *Ubc9^fKO^* and *Ubc9^fl/fl^* littermate controls (n=5,6/group). (H) Reproductive hormone levels (anti-mullerian hormone, progesterone, follicle stimulating hormone, and luteinizing hormone) measured from serum in 2-months and 6-months-old mice (n=5,7/group). Data are mean ± SEM. *p<0.05, **p<0.01 two-way ANOVA followed by Fisher’s LSD post-hoc test and unpaired two-tailed Student’s *t*-test for litter size, pup survival, and superovulation.

Our data suggest that metabolic demands of reproduction in *Ubc9^fKO^*females might interfere with reproductive function. To confirm that irregular estrous cycles imparted to *Ubc9^fKO^*subfertility, we also conducted superovulation studies. A decrease in the number of MII oocytes retrieved from young *Ubc9^fKO^* females (Figure 4G) indicates compromised ovulatory capacity. Further supporting this notion, both two- and six-month-old *Ubc9^fKO^* mice exhibited lowered levels of serum anti-Müllerian hormone (AMH) (Figure 4H), a well-established marker of the ovarian reserve that declines during aging (Buratini et al. 2022), indicating defects related to ovarian follicle dynamics. No difference in serum progesterone (P4) and follicle-stimulating hormone (FSH) were seen between genotypes. Interestingly, luteinizing hormone (LH) was higher in aged *Ubc9^fKO^* females but not littermate controls, another marker of ovarian dysfunction (Chakravarti et al. 1976). Together, the confluence of metabolic phenotyping and fertility assessments point to compromised fecundity in *Ubc9^fKO^*females.

### High-fat diet feeding normalizes estrous cycles in ‘fat-less’ mice

A sustained high-fat diet (HFD) is shown to decrease sympathetic tone, thermogenic genes, and beige fat abundance (Jun et al. 2020). To determine if we could decrease energy expenditure and beige fat activity through dietary manipulation, *Ubc9^fKO^* females and *Ubc9^fl/fl^* littermate controls were placed on HFD for 12 weeks starting at 5 weeks of age. Interestingly, *Ubc9^fKO^* females maintained similar weight gain with controls at the beginning of HFD feeding but weighed significantly less than controls after six weeks of HFD (Figure 5A). Further, CLAMS studies of *Ubc9^fKO^* females demonstrated that HFD feeding effectively normalized the previously observed hypermetabolic phenotype to levels observed in *Ubc9^fl/fl^*controls. By normalizing energy expenditure (Figure 5B) and the respiratory exchange ratio (Figure 5C), female *Ubc9^fKO^* mice on HFD were comparable to controls despite still having increased lean mass (Figure 5D). HFD feeding was also unable to rescue fat mass in these mice. Consequently, HFD caused demonstrably worsened insulin resistance (Figure 5E, ITT), further confirmed by ITT. Surprisingly, tolerance to glucose (Figure 5E, GTT) was better in female knockouts compared to control mice on HFD, likely due to high baseline insulin levels (Figure 5F). As expected, serum insulin and leptin were raised in response to HFD in *Ubc9^fl/fl^* littermate controls (Figure 5F and 5G). However, only serum insulin was increased in *Ubc9^fKO^* mice fed HFD, while leptin levels were not different compared to knockouts on normal chow (NC) (Figure 5G).

**Figure 5.**
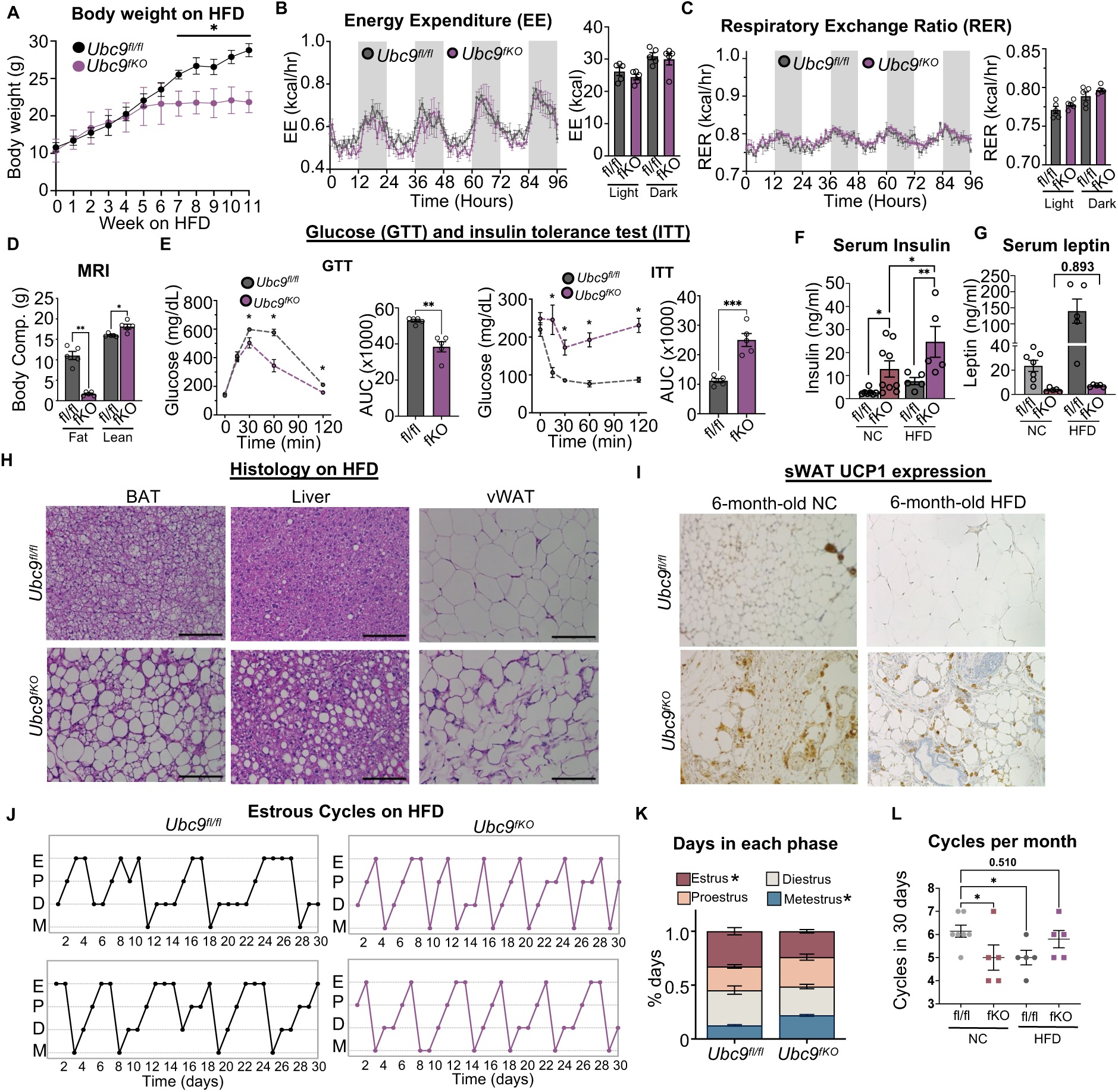
High-fat diet feeding normalizes estrous cycles in ‘fat-less’ mice. (A) Body weight measurements (g) over 11 weeks on a high-fat diet in *Ubc9^fKO^* and *Ubc9^fl/fl^* littermate controls (n=5,7/group) starting at 5 weeks of age. (B) CLAMS cages during a 96 h period from *Ubc9^fKO^* and *Ubc9^fl/fl^* on sustained HFD. Statistical analysis was performed by ANCOVA with lean body mass as a covariate for body mass-dependent variables (EE) and ANOVA for mass-independent variables (RER) (n=5/group). Data are mean ± SEM. *p<0.05, **p<0.01, ***p<0.001. (C) Respiratory exchange ratio (RER). (D) Body composition measured by Echo MRI, shown as a percentage of body weight. (E) Glucose (GTT) and insulin (ITT) tolerance test with area-under-curve (AUC) measurements from HFD-fed mice (n=5). (F) Serum insulin levels measured in normal chow (NC) and HFD-fed *Ubc9^fKO^* and age-matched *Ubc9^fl/fl^* controls. (G) Serum leptin levels measured in NC and HFD-fed *Ubc9^fKO^* and age-matched *Ubc9^fl/fl^* controls. (H) Representative images of brown adipose (BAT), liver, visceral white adipose tissue (vWAT) H&E stained histology sections from *Ubc9^fKO^* and *Ubc9^fl/fl^* mice after HFD (n=3,4/group). Images taken at 20x with scale bars at 100µm. (I) IHC staining of UCP1 in subcutaneous white adipose tissue (sWAT) taken from 6-month-old female *Ubc9^fKO^* and *Ubc9^fl/fl^* mice fed normal chow (NC) or High-fat diet (HFD) with corresponding measured expression (% area of stain). All images were taken at 20x. Scale bar set at 100µm (n=2/3group). (J) Representative 30-day estrous cycle monitoring in *Ubc9^fKO^* littermate controls on HFD (n=5/group). (K) Percent days in each phase and number of days spent in estrus over a 30-day period (n=5/group). (L) Number of cycles in 30 days between *Ubc9^fKO^* and *Ubc9^fl/fl^* mice on normal chow compared to HFD. Data are mean ± SEM. *p<0.05. **p<0.01 by two-way ANOVA followed by Fisher’s LSD post-hoc, repeated measures ANOVA for GTT and ITT, and unpaired two-tailed student’s *t*-test for AUC.

The inability of WAT depots to sequester lipids, as seen in lipodystrophy and obesity phenotypes, can cause ectopic accumulation of energy in peripheral organs, often in a sexually dimorphic manner (Lefebvre and Staels 2021; Polyzos et al. 2019). In *Ubc9^fKO^*female mice, HFD caused profound infiltration of large fat droplets in the liver and brown adipose tissue depots (Figure 5H). These data demonstrated that the additional HFD amplifies severe metabolic dysfunction in *Ubc9^fKO^*‘fat-less’ mice. To further determine if HFD-associated metabolic dysfunction was in part caused by the loss of protected compensatory beige fat, we further analyzed histological sections from sWAT. Consistent with normalized energy expenditure profiles, sWAT UCP1 in *Ubc9^fKO^* and *Ubc9^fl/fl^*is lower after HFD (Figure 5I).

Although *Ubc9^fKO^* females on HFD displayed worsened metabolic phenotypes, we wanted to see if lowered energy expenditure and loss of beige fat affected the estrous cycle irregularity in these mice. Remarkably, estrous cycles of *Ubc9^fKO^* were normalized, with proportional time spent in each phase and overall cycle regularity following the HFD intervention (Figures 5J and 5K). Countless other studies have shown that HFD feeding can result in cycle irregularity in WT mice (Liu et al. 2024). Similarly, *Ubc9^fl/fl^* controls on HFD also experienced cycle irregularity and disrupted estrous cycles. Because of the cycle irregularity displayed in control mice on HFD, we also monitored estrous cycles in age-matched mice on NC. HFD-fed *Ubc9^fKO^* mice had a comparable number of cycles per 30 days to age-matched *Ubc9^fl/fl^* mice on NC when compared by diet and genotype (Figure 5L). These data provide strong evidence that beige fat recruitment negatively impacts cycle regularity in *Ubc9^fKO^* mice.

## Discussion

In this study, we uncovered a significant trade-off between metabolic adaptations to fat loss and female reproductive function by using our previously established mouse model of lipodystrophy as a tool to study how female-specific beige fat activation impacted fertility outcomes in these mice. We found evidence of increased WAT thermogenesis in female *Ubc9^fKO^* mice, demonstrated by increases in UCP1 in sWAT and not BAT, along with other thermogenic genes and a systemic increase in energy expenditure. Prior studies demonstrated that increased thermogenesis can sustain metabolic health, despite reduced fat stores, by enhancing mitochondrial function and fatty acid utilization (Savage et al. 2005). This metabolic resilience may explain why *Ubc9^fKO^* females maintained insulin sensitivity and glucose homeostasis despite lower adiposity. Although these adaptations may respond to inadequate energy stores associated with fat loss, they appear to come at the cost of reproductive function, exhibited by disrupted estrous cycles, reduced ovarian reserve, and diminished fertility. These results also support the notion that females are particularly vulnerable to metabolic-reproductive trade-offs resulting from fat loss, as shown in other studies in which female reproduction is often halted in response to metabolic stress, nutritional availability, and famine (Della Torre and Maggi 2017; Torre et al. 2014). Importantly, to our knowledge, this is the first study to demonstrate the negative pressure beige fat imposes on female fecundity. Previous studies defined the volume of beige fat as a marker of insulin sensitivity and inversely correlate against obesity-associated metabolic dysfunction (Chouchani and Kajimura 2019; Becher et al. 2021). However, metabolic and reproductive pathways are tightly connected, and reproductive health should be considered in conjunction with metabolic health. Moreover, females have high energetic demands for reproduction, and a diversion of wasted energy to thermogenesis may decrease the energy available for reproductive function.

One important consequence of the metabolic shift in *Ubc9^fKO^* females was the disruption of estrous cycle regularity and diminished fertility. However, the observed prolongation of the estrus phase and cycle irregularity was rescued with HFD feeding. A key question arising from these findings is how cycle regularity in *Ubc9^fKO^* females is rescued with HFD feeding. Our results suggest that HFD both normalized energy expenditure and rescues estrous cycle regularity, reinforcing the idea that reproductive function is tightly linked to energy balance. However, the precise mechanism behind this rescue remains unknown. Similar to other lipodystrophic mouse models, *Ubc9^fKO^* females have lowered serum leptin, which has a well-established role in female fertility. As such, leptin add-back has been shown to rescue fertility in mouse models with fat loss and leptin deficiency (Eifler et al. 2018). Yet, it is important to note that our HFD intervention normalized estrous cycles in our model without prominently raising leptin levels, providing evidence that these are leptin-independent mechanisms.

Furthermore, our dietary intervention results further highlight the balance between energy balance, fertility, and metabolic health. While HFD improved estrous cyclicity, presumably by lowering EE and thermogenic activity, the resulting insulin resistance demonstrates a cost to metabolic function. As such, HFD may provide short-term reproductive benefits for *Ubc9^fKO^*females but, in turn, compromise insulin action in the periphery. It is possible that prolonged metabolic stress might induce irreversible alterations in ovarian function or hypothalamic-pituitary-gonadal axis regulation, preventing complete recovery of fertility. It will now be important to determine whether early nutritional or environmental interventions prevent the long-term reproductive consequences of increased energy expenditure and lean mass in females. Importantly, these results underscore the complex interplay between metabolism and reproduction and provide insight into the evolutionary prioritization of energy allocation between survival and female reproductive success.

## Resource Availability

### Lead Contact

Further information and requests for resources should be directed to and will be fulfilled by the Lead Contact, Sean M. Hartig. All data that support the findings herein presented are available from the corresponding author upon reasonable request.

### Materials Availability

Resources (*Ubc9^fKO^*mice) are available from the Lead Contact upon reasonable request.

### Data and Code Availability

All data generated or analyzed during this study are included in the published article. RNA-seq datasets are deposited in Gene Expression Omnibus.

## Acknowledgments

This work was funded by NIH R01DK114356 (S.M.H.), R01DK138018 (S.M.H.), and R01DK139397 (S.M.H.). E.S.A. received support from an American Heart Association Predoctoral Fellowship (25PRE1373489). W.D. was supported by NIH fellowship F31 AG080998. BCM core services received support from NCI P30CA125123: Genetically Engineered Rodent Models, Human Tissue Acquisition and Pathology, and Integrated Microscopy. Services and instruments used in this project for RNAseq were graciously supported, in part, by the University of Pittsburgh, the ORice of the Senior Vice Chancellor for Health Sciences, the Department of Pediatrics, the Institute for Precision Medicine, and the Richard K Mellon Foundation for Pediatric Research.

## Author Contributions

E.S.A., S.A.P., and S.M.H. conceptualized the study. A.R.C, P.K.S., and the remaining authors performed critical experiments in mice and/or contributed to the interpretation of metabolic phenotypes and biochemical outcomes. W.D. and B.Z. performed RNA-Seq analysis. E.S.A. and S.M.H. wrote the manuscript with editorial input from all authors. S.M.H. is the guarantor of this work and, as such, has full access to all the data in the study and takes responsibility for the integrity of the data and the accuracy of the data analysis.

## Declaration of generative AI and AI-assisted technologies

The authors declare no use of AI.

## Inclusion and Diversity

We support inclusive, diverse, and equitable conduct of research.

## Methods

### Animal studies

All animal procedures were approved by the Institutional Animal Care and Use Committee of Baylor College of Medicine (Animal Protocol AN-6411). Experimental animals received humane care according to criteria in the “Guide for the Care and Use of Laboratory Animals” (8th edition, revised 2011). Experimental animals were housed (no more than four per cage) in a barrier-specific pathogen-free animal facility with a 12 h dark-light cycle and free access to water and normal chow (Harlan Laboratories 2920X) unless otherwise specified. *Ubc9^fl/fl^* mice used in this study were previously generated and crossed with *Adipoq-Cre* (Jackson Laboratory #028020) to generate adipocyte-specific *Ubc9* knockout (*Ubc9^fKO^*) and littermate controls (*Ubc9^fl/fl^*) (Cox et al. 2021). Experiments were conducted using mice maintained on a C57BL/6J background. At the end of each experiment, mice were euthanized by cervical dislocation while under isoflurane anesthesia. After euthanasia, tissues were collected and either fixed in 10% formalin or flash-frozen in liquid N2 and stored at −80 °C until use. All experiments adhered to ARRIVE Guidelines. *Cre* transgenic mice were genotyped according to protocols provided by the Jackson Laboratory.

### RNA isolation and qPCR

Total RNA was extracted using the RNeasy Lipid Tissue kit (Qiagen). cDNA was synthesized from total RNA using SuperScript IV VILO Master Mix (Invitrogen # 11766050). Relative mRNA expression was measured with SsoAdvanced Universal Probes Supermix reactions (Bio-Rad #175284) read out with a QuantStudio 3 real-time PCR system (Applied Biosystems). TATA-box binding protein (Tbp) was the invariant control. Roche Universal Probe Gene Expression Assays (mouse) or individual TaqMan Gene Expression Assays (Thermo Fisher) were used as previously described (Cox et al. 2022).

### Indirect calorimetry

*Ubc9*^fKO^ mice and littermate controls (*Ubc9*^fl/fl^) were maintained on normal chow and housed at room temperature in Comprehensive Lab Animal Monitoring System (CLAMS) home cages (Columbus Instruments). Oxygen consumption (VO2), carbon dioxide production (VCO2), energy expenditure (EE), food and water intake, and activity were measured for six days (BCM Mouse Metabolic and Phenotyping Core). Mouse body weight was recorded, and body composition was measured by magnetic resonance imaging (Echo Medical Systems) prior to indirect calorimetry. The first two days (48 hr) of measurements were excluded from the analysis to allow for the acclamation of CLAMS cages. Statistical analysis of mass-dependent variables (VO2, VCO2, EE, and food and water intake) was performed by ANCOVA with lean body mass as a co-variate and ANOVA for mass-independent variables (wheel running, respiratory exchange ratio, and physical activity) using the CalR web-based tool (Mina et al. 2018).

### Glucose and insulin tolerance tests

To assess glucose tolerance, mice were fasted for 16 h, and glucose (1.5 g/kg body weight) was administered by intraperitoneal (IP) injection. To assess insulin tolerance, mice were fasted four hours prior to insulin IP injection (1.5 U/kg body weight). Blood glucose levels were measured by handheld glucometer.

### ELISAs and FFA assays

Serum collected from ad-libitum-fed mice was used to measure insulin (Millipore #EZRMI-13K), leptin (Crystal Chem #90030), adiponectin (Thermo Fisher #KMP0041), and free fatty acids (ZenBio #sfa-1). The serum was also analyzed for triglycerides (Thermo Fisher TR22421) and total cholesterol (Thermo Fisher TR13421).

### Histology

Formalin-fixed, paraffin-embedded adipose sections and liver were stained with hematoxylin and eosin (H&E) by the BCM Human Tissue Acquisition and Pathology Core (HTAP). Images were captured (20X) using a Nikon Ci-L Brightfield microscope.

### Immunohistochemistry

Immunohistochemistry was performed on formalin-fixed, paraffin-embedded adipose tissue sections by the BCM Human Tissue Acquisition and Pathology Core for beige adipocyte marker UCP1 (abcam ab10983).

### RNA sequencing

Tissue was weighed, and 30-50mg of tissue was used for RNA extraction. RNA was extracted using an RNeasy Lipid Tissue kit (Qiagen: 74804) along with a Rnase-free DNase set (Qiagen: 79254) for DNA digestion. Library generation and sequencing were performed in the Health Sciences Sequencing Core at UPMC Children’s Hospital of Pittsburgh, Rangos Research Center. RNA was assessed for quality using an Agilent TapeStation 4150/Fragment Analyzer 5300, and RNA concentration was quantified on a Qubit FLEX fluorometer. Libraries were generated with the Illumina Stranded Total Library Prep kit (Illumina: 20040529) according to the manufacturer’s instructions. Library quantification and assessment were done using a Qubit FLEX fluorometer and an Agilent TapeStation 4150/Fragment Analyzer 5300. Sequencing was performed on an Illumina NextSeq 2000, using a P3 flow cell with read lengths of 2×101 bp, with a target of 80 million reads per sample. Sequencing data was demultiplexed by the on-board Illumina DRAGEN FASTQ Generation software (v3.10.12). Bioinformatics work was done through Galaxy (Blankenberg et al., 2010; Ostrovsky et al., 2022). Raw reads were processed with *Trim Galore!* (Krueger, 2015), alignment was performed with *RNA STAR* (Dobin et al., 2013) using the mm10 reference genome, reads were counted with *htseq-count* (Anders et al., 2015), FPKM values were determined with *Cu9links* (Trapnell et al., 2010), and diRerential gene expression analysis was performed with *DESeq2* (Love et al., 2014). Volcano plots were generated using VolcaNoseR (Goedhart and Luijsterburg, 2020). Venn diagrams were generated using DeepVenn (Hulsen, 2022), and DAVID was used to determine KEGG_PATHWAY enrichment (Kanehisa and Goto, 2000; Sherman et al., 2022) in upregulated genes (log2 fold change greater than or equal to 1 and adjusted p values less than 0.05). Data is available through the NCBI Gene Expression Omnibus (GSE289065).

### Fertility analysis

To test fecundity, sexually mature (8-week-old) *Ubc9^fKO^*females or control littermates were pair-housed with sexually mature (8-week-old) C57BL/6J wild-type males and continuously mated for 6 months. An n=10/group started the fertility study, but only mice n=8/group survived to completion and were analyzed due to pregnancy complications resulting in animal welfare concerns. Females were monitored daily for new litters, and the number of pups, date of birth, and sex were recorded. Any resulting litters were weaned and weighed at 3 weeks of age. For estrous cycle monitoring, mice were individually housed, and vaginal lavage and cytology were performed daily for 1 month as described(Byers et al. 2012).

### Superovulation experiments

Female mice aged 5 weeks were injected with 5 IU pregnant mare serum gonadotropin (PMSG; ProSpecBio), and 46 h later injected with 5 IU human chorionic gonadotropin (hCG; Pregnyl; Merck Pharmaceuticals) by intraperitoneal (IP) injection. Mice were euthanized 16-18 h after injection with hCG, and cumulus-oocyte complexes were harvested from the oviduct ampulla and incubated with 0.3 mg/ml hyaluronidase (Sigma-Aldrich) to detach cumulus cells and MII oocytes counted.

### Reproductive hormone analysis

Blood was collected via cardiac puncture from isoflurane-anesthetized group-housed mice, and the serum was separated by centrifugation in microtainer collection tubes (Becton, Dickinson, and Company) and frozen at −20°C until assayed. Mouse LH, FSH, AMH, and progesterone levels were analyzed in phase-matched mice via ELISA at the University of Virginia Ligand Core Facility [Specialized Cooperative Centers Program in Reproductive Research, National Institute of Child Health and Human Development (NICHD)/National Institutes of Health (NIH) U54-HD28934]. The assay method information is available online (https://med.virginia.edu/research-in-reproduction/ligand-assay-analysis-core/assay-methods/).

### Statistical analysis

Unless otherwise noted, statistical analyses were performed using GraphPad Prism GraphPad Prism 9 (GraphPad Software, La Jolla, CA). A two-tailed unpaired Student t-test was used for single comparisons. One-way analysis of variance followed by Fisher’s least significant difference test was used for multiple comparisons. For gene expression data, statistical significance was assessed by multiple unpaired t-tests with a q-value < 0.05. Statistical analysis of energy balance was performed by ANCOVA with lean body mass as a co-variate and cumulative food intake by standard ANOVA using the CalR web-based tool(Mina et al. 2018).

A power analysis was performed for all experimental methods, and sample sizes are indicated in the text and figure legends. All data are presented as mean ± standard error of the mean (SEM). Statistical significance shown as * p < 0.05, ** p < 0.01, *** p < 0.001, *** p < 0.0001.

## References

Agarwal N, Iyer D, Saha P, Cox AR, Xia Y, Utay NS, Somasundaram A, Schubert U, Lake JE, Hartig SM, et al. 2021. HIV-1 Viral Protein R Couples Metabolic Inflexibility With White Adipose Tissue Thermogenesis. Diabetes 70: 2014–2025.

Becher T, Palanisamy S, Kramer DJ, Eljalby M, Marx SJ, Wibmer AG, Butler SD, Jiang CS, Vaughan R, Schöder H, et al. 2021. Brown adipose tissue is associated with cardiometabolic health. Nat Med 27: 58–65.

Béréziat V, Cervera P, Le Dour C, Verpont M-C, Dumont S, Vantyghem M-C, Capeau J, Vigouroux C, Lipodystrophy Study Group. 2011. LMNA mutations induce a non-inflammatory fibrosis and a brown fat-like dystrophy of enlarged cervical adipose tissue. Am J Pathol 179: 2443–2453.

Boutari C, Pappas PD, Mintziori G, Nigdelis MP, Athanasiadis L, Goulis DG, Mantzoros CS. 2020. The effect of underweight on female and male reproduction. Metabolism 107: 154229.

Buratini J, Dellaqua TT, Dal Canto M, La Marca A, Carone D, Mignini Renzini M, Webb R. 2022. The putative roles of FSH and AMH in the regulation of oocyte developmental competence: from fertility prognosis to mechanisms underlying age-related subfertility. Hum Reprod Update 28: 232–254.

Byers SL, Wiles MV, Dunn SL, Taft RA. 2012. Mouse Estrous Cycle Identification Tool and Images. PLOS ONE 7: e35538.

Chakravarti S, Collins WP, Forecast JD, Newton JR, Oram DH, Studd JW. 1976. Hormonal profiles after the menopause. Br Med J 2: 784–787.

Chen Y, Ikeda K, Yoneshiro T, Scaramozza A, Tajima K, Wang Q, Kim K, Shinoda K, Sponton CH, Brown Z, et al. 2019. Thermal stress induces glycolytic beige fat formation via a myogenic state. Nature 565: 180–185.

Chouchani ET, Kajimura S. 2019. Metabolic adaptation and maladaptation in adipose tissue. Nat Metab 1: 189–200.

Cox AR, Chernis N, Kim KH, Masschelin PM, Saha PK, Briley SM, Sharp R, Li X, Felix JB, Sun Z, et al. 2021. Ube2i deletion in adipocytes causes lipoatrophy in mice. Mol Metab 48: 101221.

Cox AR, Masschelin PM, Saha PK, Felix JB, Sharp R, Lian Z, Xia Y, Chernis N, Bader DA, Kim KH, et al. 2022. The rheumatoid arthritis drug auranofin lowers leptin levels and exerts anti-diabetic effects in obese mice. Cell Metab 34: 1932–1946.e7.

Cypess AM, Lehman S, Williams G, Tal I, Rodman D, Goldfine AB, Kuo FC, Palmer EL, Tseng Y-H, Doria A, et al. 2009. Identification and Importance of Brown Adipose Tissue in Adult Humans. N Engl J Med 360: 1509–1517.

Della Torre S, Maggi A. 2017. Sex Differences: A Resultant of an Evolutionary Pressure? Cell Metabolism 25: 499–505.

Eifler L, Hoffmann A, Wagner IV, Klöting N, Sahlin L, Ebert T, Jessnitzer B, Lössner U, Stumvoll M, Söder O, et al. 2018. Leptin restores markers of female fertility in lipodystrophy. Biochimica et Biophysica Acta (BBA) - Molecular Basis of Disease 1864: 3292–3297.

Gallego-Escuredo JM, Domingo P, Fontdevila J, Villarroya J, Domingo JC, Martinez E, Giralt M, Villarroya F. 2013. Hypertrophied facial fat in an HIV-1-infected patient after autologous transplantation from “buffalo hump” retains a partial brown-fat-like molecular signature. Antivir Ther 18: 635–639.

Ghaben AL, Scherer PE. 2019. Adipogenesis and metabolic health. Nat Rev Mol Cell Biol 20: 242–258.

Guo B, Liu J, Wang B, Zhang C, Su Z, Zhao M, Qin L, Zhang W, Zheng R. 2022. Withaferin A Promotes White Adipose Browning and Prevents Obesity Through Sympathetic Nerve-Activated Prdm16-FATP1 Axis. Diabetes 71: 249–263.

Jakobsson A, Jörgensen JA, Jacobsson A. 2005. Differential regulation of fatty acid elongation enzymes in brown adipocytes implies a unique role for Elovl3 during increased fatty acid oxidation. American Journal of Physiology-Endocrinology and Metabolism 289: E517–E526.

Jun H, Ma Y, Chen Y, Gong J, Liu S, Wang J, Knights AJ, Qiao X, Emont MP, Xu XZS, et al. 2020. Adrenergic-independent signaling via CHRNA2 regulates beige fat activation. Dev Cell 54: 106–116.e5.

Kim S-N, Jung Y-S, Kwon H-J, Seong JK, Granneman JG, Lee Y-H. 2016. Sex differences in sympathetic innervation and browning of white adipose tissue of mice. Biol Sex Differ 7: 67.

Labbé SM, Caron A, Chechi K, Laplante M, Lecomte R, Richard D. 2016. Metabolic activity of brown, “beige,” and white adipose tissues in response to chronic adrenergic stimulation in male mice. Am J Physiol Endocrinol Metab 311: E260–268.

Lefebvre P, Staels B. 2021. Hepatic sexual dimorphism — implications for non-alcoholic fatty liver disease. Nat Rev Endocrinol 17: 662–670.

Liu C, Dou Y, Zhang M, Han S, Hu S, Li Y, Yu Z, Liu Y, Liang X, Chen Z-J, et al. 2024. High-fat and high-sucrose diet impairs female reproduction by altering ovarian transcriptomic and metabolic signatures. J Transl Med 22: 145.

Mauvais-Jarvis F. 2024. Sex differences in energy metabolism: natural selection, mechanisms and consequences. Nat Rev Nephrol 20: 56–69.

Mina AI, LeClair RA, LeClair KB, Cohen DE, Lantier L, Banks AS. 2018. CalR: A Web-Based Analysis Tool for Indirect Calorimetry Experiments. Cell Metab 28: 656–666.e1.

Pellegrini C, Columbaro M, Schena E, Prencipe S, Andrenacci D, Iozzo P, Angela Guzzardi M, Capanni C, Mattioli E, Loi M, et al. 2019. Altered adipocyte differentiation and unbalanced autophagy in type 2 Familial Partial Lipodystrophy: an in vitro and in vivo study of adipose tissue browning. Exp Mol Med 51: 1–17.

Petruzzelli M, Schweiger M, Schreiber R, Campos-Olivas R, Tsoli M, Allen J, Swarbrick M, Rose-John S, Rincon M, Robertson G, et al. 2014. A Switch from White to Brown Fat Increases Energy Expenditure in Cancer-Associated Cachexia. Cell Metabolism 20: 433–447.

Polyzos SA, Perakakis N, Mantzoros CS. 2019. Fatty liver in lipodystrophy: A review with a focus on therapeutic perspectives of adiponectin and/or leptin replacement. Metabolism - Clinical and Experimental 96: 66–82.

Rodríguez de la Concepción ML, Domingo JC, Domingo P, Giralt M, Villarroya F. 2004. Uncoupling protein 1 gene expression implicates brown adipocytes in highly active antiretroviral therapy-associated lipomatosis. AIDS 18: 959–960.

Santos RS, Frank AP, Fátima LA, Palmer BF, Öz OK, Clegg DJ. 2018. Activation of estrogen receptor alpha induces beiging of adipocytes. Molecular Metabolism 18: 51–59.

Savage DB, Murgatroyd PR, Chatterjee VK, O’Rahilly S. 2005. Energy Expenditure and Adaptive Responses to an Acute Hypercaloric Fat Load in Humans with Lipodystrophy. The Journal of Clinical Endocrinology & Metabolism 90: 1446–1452.

Srinivasa S, Torriani M, Fitch KV, Maehler P, Iyengar S, Feldpausch M, Cypess AM, Grinspoon SK. 2019. Brief Report: Adipogenic Expression of Brown Fat Genes in HIV and HIV-Related Parameters. J Acquir Immune Defic Syndr 82: 491–495.

Torre SD, Benedusi V, Fontana R, Maggi A. 2014. Energy metabolism and fertility—a balance preserved for female health. Nat Rev Endocrinol 10: 13–23.

Wang F, Mullican SE, DiSpirito JR, Peed LC, Lazar MA. 2013. Lipoatrophy and severe metabolic disturbance in mice with fat-specific deletion of PPARγ. Proceedings of the National Academy of Sciences 110: 18656–18661.

Westerberg R, Månsson J-E, Golozoubova V, Shabalina IG, Backlund EC, Tvrdik P, Retterstøl K, Capecchi MR, Jacobsson A. 2006. ELOVL3 is an important component for early onset of lipid recruitment in brown adipose tissue. J Biol Chem 281: 4958–4968.

Zhu L, Zhou B, Zhu X, Cheng F, Pan Y, Zhou Y, Wu Y, Xu Q. 2022. Association Between Body Mass Index and Female Infertility in the United States: Data from National Health and Nutrition Examination Survey 2013–2018. Int J Gen Med 15: 1821–1831.

